# A Framework for Systematically Evaluating the Representations Learned by A Deep Learning Classifier from Raw Multi-Channel Electroencephalogram Data

**DOI:** 10.1101/2023.03.20.533467

**Authors:** Charles A. Ellis, Abhinav Sattiraju, Robyn L. Miller, Vince D. Calhoun

## Abstract

The application of deep learning methods to raw electroencephalogram (EEG) data is growing increasingly common. While these methods offer the possibility of improved performance relative to other approaches applied to manually engineered features, they also present the problem of reduced explainability. As such, a number of studies have sought to provide explainability methods uniquely adapted to the domain of deep learning-based raw EEG classification. In this study, we present a taxonomy of those methods, identifying existing approaches that provide insight into spatial, spectral, and temporal features. We then present a novel framework consisting of a series of explainability approaches for insight into classifiers trained on raw EEG data. Our framework provides spatial, spectral, and temporal explanations similar to existing approaches. However, it also, to the best of our knowledge, proposes the first explainability approaches for insight into spatial and spatio-spectral interactions in EEG. This is particularly important given the frequent use and well-characterized importance of EEG connectivity measures for neurological and neuropsychiatric disorder analysis. We demonstrate our proposed framework within the context of automated major depressive disorder (MDD) diagnosis, training a high performing one-dimensional convolutional neural network with a robust cross-validation approach on a publicly available dataset. We identify interactions between frontal and central electrodes and other electrodes and identify differences in frontal δ, θ, β, and γ_low_ between healthy controls and individuals with MDD. Our study represents a significant step forward for the field of deep learning-based raw EEG classification, providing new capabilities in interaction explainability and providing direction for future innovations through our proposed taxonomy.

## INTRODUCTION

In recent years, studies have increasingly applied deep learning approaches to raw electroencephalography (EEG) data. Relative to studies using traditional machine learning and deep learning methods with extracted features, deep learning studies using raw EEG allow for automated feature learning and the discovery of EEG features that might ordinarily be overlooked. This benefit has the potential to enhance model performance. Nevertheless, the use of raw EEG also occasions an important shortcoming. Namely, deep learning models with raw EEG are not as explainable as traditional machine learning [1] or deep learning models [2], [3] applied to extracted features. This has resulted in the development of a subfield of EEG analysis seeking to make deep learning models with raw EEG more explainable. To a large extent, these studies have succeeded. Under certain circumstances, EEG explainability approaches can provide insight into key channels [4]–[11], frequency bands [4], [5], [8], [12]–[18], and waveforms [5], [8], [14], [16], [17]. However, existing methods do not provide insight into interactions between channels. In this study, we present a taxonomy of deep learning-based raw EEG explainability approaches, identifying critical gaps in the capabilities of the field. We then present a series of explainability approaches that form a framework for systematically evaluating what a deep learning model has learned from raw EEG. Specifically, we train a high-performing one-dimensional convolutional neural network (1D-CNN) with a robust cross-validation approach to differentiate between healthy individuals and individuals with clinically diagnosed major depressive disorder (MDD) on multichannel EEG data. We present approaches to (1) identify the relative importance of each channel, (2) identify interactions uncovered by the model between channels, (3) identify key frequency bands in each channel, (4) identify interactions between frequency bands in each channel and other channels, and (5) identify representative samples of each class and the waveforms of importance to their classification. Our identification of spatio-spectral interactions is to our knowledge the first implementation of such a method in raw EEG-based deep learning explainability. Moreover, our study represents a significant step forward for the domain of raw EEG-based deep learning explainability and has the potential to stimulate future advances in the field.

### Modalities Used for Analysis of Neurological and Neuropsychiatric Disorders and Advantage of EEG

Multiple modalities have been used to study neurological and neuropsychiatric disorders. A few of these modalities include EEG [1]–[3], [9], [10], [19]–[27], magnetoencephalography (MEG) [28]–[30], and functional magnetic resonance imaging (fMRI) [30]–[37]. Each modality offers both advantages and disadvantages. For example, fMRI has enhanced spatial resolution relative to EEG and MEG. However, EEG and MEG have significantly improved temporal resolution relative to fMRI, which can afford better insight into the effects of disorders upon brain dynamics. Additionally, relative to MEG, EEG devices can be performed much more cheaply and are more widespread, making it better suited for deployment in a clinical setting. Within the domain of EEG analysis, both task [8], [11], [38] and resting-state [5], [6], [9], [10], [25], [39], [40] analyses are commonly performed. However, most brain activity spontaneously occurs (i.e., reflects brain networks that are unmodulated by any task), so resting-state activity better reflects the activity that is common to an individual with a disorder. Additionally, individuals with a disorder may perform a task less effectively than healthy individuals, which could introduce a confounder in any subsequent analyses [41]. As such, in this study, we focus on explainability for resting-state EEG.

### Features Commonly Included in EEG Analyses

Historically, many features have been extracted from EEG for insight into neurological and neuropsychiatric disorders. These include single-channel features like spectral power [1]–[3], [18], [25], [28], [42]–[45] and multi-channel features like temporal and spectral connectivity [26], [27], [46]–[49]. Spectral power has been associated with disorders like schizophrenia [43], attention deficit hyperactivity disorder (ADHD) [43], obsessive compulsive disorder (OCD) [43], Parkinson’s disease, and major depressive disorder [50]. Connectivity features have shown high discriminative power and been associated with a many disorders including schizophrenia [49], MDD [27], and Alzheimer’s disease [51]. Importantly, previous studies of MDD have identified effects upon connectivity between all frequency bands [27] and between frontal electrodes, between temporal electrodes, and between temporal and central electrodes [26].

### Transition from Manually Engineered Features to Raw EEG Data

Building upon these features, many studies have trained machine learning [25], [28], [42] and deep learning [2], [3], [18], [19], [39], [45], [52] models on spectral power features. Additionally, a few studies have trained machine learning [47] and deep learning models [48] on extracted connectivity features. These studies have obtained high levels of model performance while simultaneously offering high levels of explainability. They have obtained high levels of explainability largely because many methods have been previously developed to explain traditional machine learning models [53]–[55] and many methods like saliency [56], gradient-weighted class activation mapping (Grad-CAM) [57], and layer-wise relevance propagation (LRP) [58] have been developed within the domain of image classification to explain deep learning models. However, models using extracted features have an inherit limitation. Namely, they restrict the space of features over which models can learn. As such, over time, as the field of deep learning has further developed, an increasing number of studies have begun training deep learning models on raw EEG data [4]–[14], [16]–[18]. Deep learning models employ an automated feature extraction approach that precludes the need for manually engineered features. As such, deep learning models are theoretically able to learn from the entire feature space when applied to raw EEG data. Unfortunately, they also have reduced explainability due to the high dimensionality of the input data.

### Explainability in Models with Manually Engineered Versus Automatically Learned Features

Deep learning models applied to raw EEG are not less explainable because existing explainability methods cannot be applied in the context of EEG. Rather, deep learning models applied to raw EEG are less explainable because the temporal nature of EEG data presents unique problems relative to tabular and image data. Traditional explainability methods cannot be directly translated to provide insight into key frequency bands because the input to models is a time-series. Traditional methods [59] cannot be directly translated to identify key waveforms because to extract useful global insight thousands or hundreds of thousands of samples might need to be analyzed. Traditional methods [60] that account for interactions are also difficult to translate directly to EEG because of the large number of features per sample of EEG data (e.g., a sample may have 19 to 60 channels and be thousands of time points long). As such, over time a growing number of studies have begun seeking to develop explainability methods uniquely adapted to the domain of deep learning-based raw EEG analysis. It should be noted, however, that methods like those developed for multimodal data explainability [6], [7], [61], [62] can be adapted to multichannel EEG data with minimal inconvenience.

### Taxonomy of Explainability Methods for Deep Learning Models Trained on Raw EEG

In this section, we describe a taxonomy of explainability methods for deep learning models trained on raw EEG. As shown in Figure 1, deep learning-based explainability methods for raw EEG can be categorized on a hierarchy with two general levels: (1) based on the traditional features into which they provide insight and (2) based on the mechanisms by which they provide that insight. Existing explainability approaches can generally provide insight into 3 types of features: (1) spatial features (i.e., identifying specific brain regions or electrodes of importance), (2) spectral features (i.e., identifying specific frequency bands of importance), and (3) temporal features (i.e., identifying specific waveforms of interest). Approaches for identifying spatial or multimodal importance typically use some variation of ablation [6], [7], [9], [14], [63], [64], in which information from a particular channel is removed and the effect upon model performance or softmax activations is quantified, or a gradient-based feature attribution (GBFA) approach [65] like LRP [61], [63], in which importance is summed across all time points for each channel.

**Figure 1.**
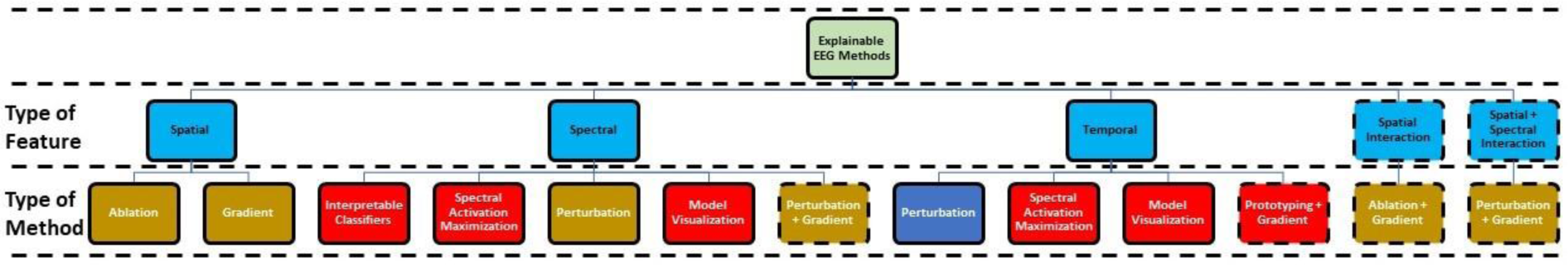
Taxonomy of Explainability Approaches for Deep Learning Models Using Raw EEG Data. The taxonomy has 3 levels that are each separated by black horizontal dashed lines: (1) the overall field of EEG explainability methods, (2) the types of features into which the explainability methods provide insight, and (3) the types of explainability approaches that provides insight into the different feature types. Light blue boxes correspond to specific types of features. Note that the type of method or explainability approach generally corresponds to methods that were first developed outside the domain of EEG analysis and then adapted to the domain. Dark blue, red, and gold boxes show methods that provide local, global, and both local and global explanations. Boxes surrounded by dashed black lines show explainability approaches first proposed in this study.

Approaches for identifying spectral importance generally fall within one of four categories. (1) They use interpretable classifiers with filters designed to extract specific frequencies [4], [66]. While highly innovative, these classifiers still inherently limit the space of possible features. (2) They use methods like activation maximization [56]. Two studies have sought to identify the frequencies at which sinusoids maximize the activation of early convolutional layers [14], [67], and one study sought to optimize the multi-spectral content of a sample to maximize the activation of the final softmax output layer in a class-specific manner [16]. These approaches do not actually indicate the importance of frequency bands to a classifier. Rather, they show a set of frequencies that are extracted at a particular layer or representative of a particular class. As such, they can produce a highly useful representation of what a model has learned. It should be noted that in methods like that in [16] there are theoretically multiple possible representations that could have high class-specific activations, so resulting samples may not contain all of the features important to the capacity of the model to identify each class. (3) A number of studies have used the fast Fourier transform to convert to and from the frequency domain wherein the perturbed specific frequency bands and examined the resulting effect upon model performance [9], [13] or predictions [12], [68]. These methods are highly effective. However, in some models, if performed on training data or sometimes even test data, perturbation may have a negligible effect upon model performance or activations, and an explanation may not be obtained. As an example, note how perturbation of frequency bands in the Awake class sample, which had an extremely high activation, had minimal effect upon the model activation in [16]. (4) Several studies have created specialized CNN architectures with extended first layer filters capable of extracting distinct waveforms [5], [40], [67]. The filters can then be converted to the frequency domain and visualized. These methods provide a very effective way for understanding what frequencies were extracted by a model. Additionally, when combined with perturbation, they can also provide an effective approach for estimating spectral importance. Nevertheless, they require the development of a highly specialized architecture and are thus incompatible with many architectures developed within the field.

Approaches for temporal waveform importance generally fall into one of three categories. Additionally, while these approaches can be effective to a degree, they all have key shortcomings, and there is generally significant room for continued innovation within this type of EEG explainability. (1) Windows of individual samples can be perturbed. Those windows that cause the largest change in softmax layer activation can then be considered important [14]. While this approach can give insight into the importance of individual waveforms for the classification of an individual sample, the perturbation of individual time windows may not always have a significant effect upon the softmax layer activation. A key shortcoming is that the identified waveforms also cannot be assumed to be of global significance, and it is impractical to perturb windows across a dataset with thousands or hundreds of thousands of samples. (2) Similar to approaches in spectral importance, activation maximization [56] can be applied to identify key waveforms [16]. Activation maximization approaches have been applied to other types of time-series classification [69], [70]. The methods in these two studies optimize the content of a sample in the time domain and are effective for short time-series (i.e., around 30 time points long). However, when applied to longer time-series like those found in resting-state EEG, they tend to do a very poor job creating recognizable waveforms [16]. This led to one study optimizing the spectral content of a sample to create waveforms [16]. This approach obtains more realistic waveforms than the approaches shown in other domains for shorter time-series [69], [70] but still leaves room for improvement. (3) Lastly, model visualization approaches that provide insights into important spectral features can also identify the importance of waveforms when paired with perturbation of model filters [5], [8], [40], [67]. However, while these approaches give the clearest insight into identified waveforms, they also require the design of special architectures that may not be able to obtain high levels of classification performance for all applications. It should also be noted that there is a tentative fourth category for identifying temporal waveform significance. GBFA methods like LRP can be applied to identify the relative importance of waveforms. This is a tentative category because while the approach has been used to identify important time points [71], it has not yet been applied to identify key waveforms. As demonstrated in several fMRI classification studies, this approach can also provide global insight into patterns of importance distribution [72], [73].

While significant advancement has been made in the field of deep learning-based explainability for raw EEG, there is still significant room for continued innovation and development. As previously described, models may sometimes not be sensitive enough to existing perturbation approaches to produce a significant change in model softmax activations or performance, which can prevent the approaches from providing usable explanations [16]. Additionally, the most effective approaches for insights into temporal waveform importance require the use of specially designed classifiers [5], [8], [40], [67], and there is a need for approaches that can be applied to any deep learning architecture. Lastly, while we have not yet mentioned this shortcoming and some explainability approaches have been applied to multichannel EEG data, many existing EEG explainability approaches have been developed within the context of single channel sleep stage classification. This is likely due to (1) the well-characterized features of sleep stages [74], (2) the comparative ease of developing models for sleep stage classification, and (3) the public availability of large EEG sleep stage datasets [75]–[77]. As such, there is a need to extend these approaches to multichannel EEG data, which is often used for more complex classification tasks [9], [21], [24], [26], [27], [78]–[81]. Related to this problem of multichannel explainability, is that, to the best of our knowledge, no existing approaches have sought to provide insight into interactions between different frequency bands and channels, which is a key limitation given the relative importance of connectivity-based features in models using traditional feature extraction [46]–[48].

In this study, we expand on the taxonomy that we previously presented. We present a systematic framework for evaluating the features of a deep learning classifier that includes inter-channel and spatio-spectral interaction - a novel feature type for deep learning-based raw EEG explainability. As such, our framework encompasses spatial, spectral, temporal, and interaction-based explanations. We (1) identify the relative importance of each channel, (2) identify interactions uncovered by the model between channels, (3) identify key frequency bands in each channel, (4) identify interactions between frequency bands in each channel and other channels, and (5) identify representative samples of each class and the waveforms of importance to their classification. We present our approach within the context of explaining a 1D-CNN trained on data from individuals with MDD (MDDs) and healthy controls (HCs). Our framework represents a significant step forward for the field of deep learning-based raw EEG explainability, and we hope that it will inspire future methods also capable of solving the problems that we presented in our taxonomy of explainability approaches.

## METHODS

In this section, we describe our proposed framework. As detailed in Figure 2, we (1) used multi-channel resting-state EEG data from 28 healthy controls (HCs) and 30 individuals with MDD (MDDs). (2) We trained a one-dimensional convolutional neural network (1D-CNN) for classification and evaluated model performance. (3) We applied layer-wise relevance propagation (LRP) to identify the relative importance of each channel, and (4) we applied a combination of ablation and LRP to identify interactions in the representations learned by the model between channels. We applied a combination of LRP and spectral perturbations to identify (5) the relative importance of each canonical frequency band in each channel and (6) interactions between the representations learned by the model for the canonical frequency bands in each channel and every other channel. (7) We applied a combination of a novel prototyping approach and LRP to identify representative samples of each class and identify important waveforms that the model used to differentiate them. Our code is publicly available on GitHub and can be found at: https://github.com/cae67/MultichannelExplainabilityFramework.git.

**Figure 2.**
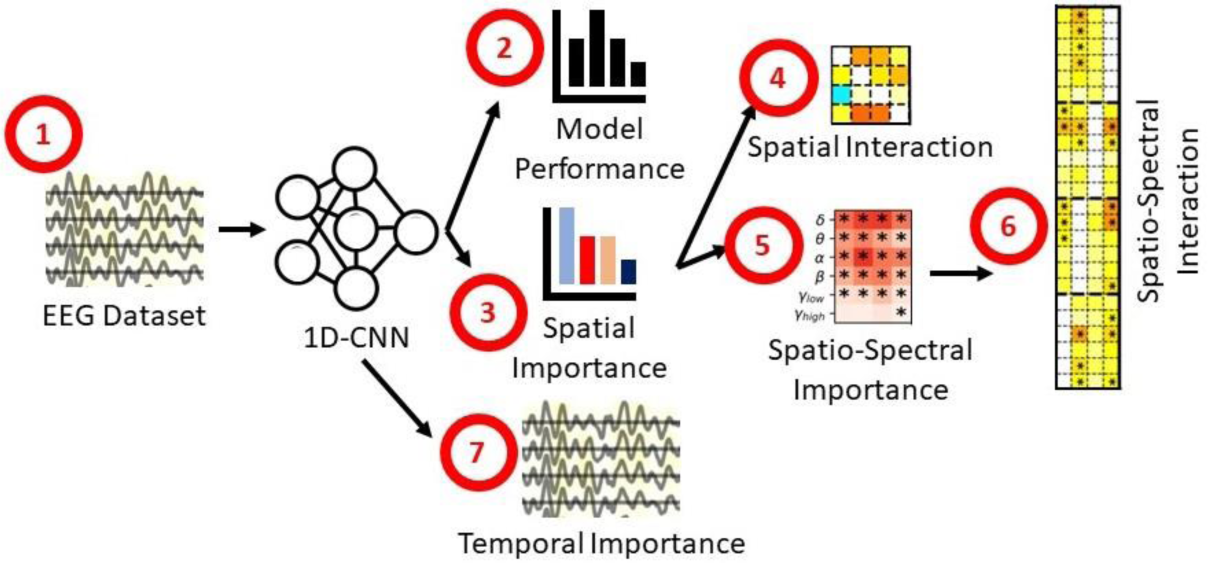
Overview of Methods. (1) We used a publicly available resting-state EEG dataset containing data from healthy individuals and individuals with major depressive disorder (MDD). (2) We first trained a one-dimensional convolutional neural network for automated MDD diagnosis and evaluated overall model performance. (3) We applied layer-wise relevance propagation (LRP) to identify the relative importance of each electrode (i.e., spatial importance). (4) We combined LRP with ablation to quantify how much the amount of LRP relevance assigned to each channel changed following the perturbation of other channels (i.e, spatial interaction). (5) We combined LRP with spectral perturbation to quantify how much the amount of LRP relevance assigned to a channel changed following the perturbation of frequency bands within that channel (i.e., spatio-spectral importance). (6) We combined LRP with spectral perturbation to quantify how much the amount of LRP relevance assigned to a channel changed following the perturbation of frequency bands within other channels (i.e., spatio-spectral importance). (7) Lastly, we combined a prototyping approach with LRP to identify representative samples of each class and to identify the relative importance of waveforms in each of those samples (i.e., temporal importance).

### Description of Data Acquisition and Preprocessing

We used a publicly available scalp EEG dataset [23] consisting of 30 MDDs and 28 age-matched HCs between the ages of 12 to 77 that has been used in multiple studies [24], [26], [78]. The data can be found at https://figshare.com/articles/dataset/EEG_Data_New/4244171. While we were not involved with data collection, we detail the collection procedures below. MDD participants met the diagnostic criteria for MDD defined in the Diagnostic and Statistical Manual-IV (DSM-IV) [82]. Common symptoms of MDD include a depressed mood, a loss of interest or pleasure, changes in appetite or weight, psychomotor agitation, feelings of worthlessness or excessive guilt, diminished ability to concentrate, and frequent thoughts of death [83]. HCs were determined to be healthy following examination for psychiatric conditions. To avoid potential confounding effects of medication, all MDDs underwent a two-week washout period prior to the first EEG recordings. All participants gave informed consent prior to data collection that was approved by the human ethics committee of the Hospital Universiti Sains Malaysia (HUSM) in Kelantan, Malaysia.

While separate recordings were performed at resting state with both eyes open and eyes closed for each participant, we only used data from recordings with eyes closed in this study. Participants were instructed to sit in a semi-recumbent position and minimize head movements and eye blinks. The Brain Master Discovery amplifier (Make: Brain Master, Model: Discovery 24e, Manufacturer: Brainmaster Technologies Inc.) was used to amplify EEG signals from the sensors. Recordings were performed for 5 to 10 minutes with a sampling rate of 256 Hertz (Hz) using a standard 10-20 format with 64 electrodes. The data were band pass filtered from 0.1 to 70 Hz and were notch filtered at 50 Hz to remove line-related noise. EEG data were recorded with the linked ear reference and were re-referenced to the infinity reference [84].

Due to the high levels of correlation present between scalp EEG channels, we only used a subset of electrodes. This approach is similar to those of other studies of neuropsychiatric disorders [9], [20], [25], [85]. Specifically, we used the Fp1, Fp2, F7, F3, Fz, F4, F8, T3, C3, Cz, C4, T4, T5, P3, Pz, P4, T6, O1, and O2 electrodes. We downsampled the data from 256 Hz to 200 Hz. To increase the number of samples available for classification, we used a 25-second sliding window with a 2.5-second sliding step size to separate the recordings into epochs. After dividing the data into epochs, we channel-wise z-scored the epochs for each participant separately. Our final dataset consisted of 2,950 SZ epochs and 2,942 HC epochs. Importantly, we did not remove any samples with extreme amplitude values.

### Description of Model Development

We adapted an architecture (Figure 3) that was originally developed in [85] for schizophrenia classification and was later used in [78] for MDD classification. We implemented the model in Keras 2.2.4 [86] to maintain compatibility with explainability libraries. Input samples had dimensions of 5,000 time points x 19 channels. Relative to [85], we added multiple batch normalization layers and converted ReLU activation functions to ELU activation functions. We also modified the training approach from [78] in an effort to enhance model performance. As mentioned in [27], many previous studies involving classification of MDD EEG data have used poor cross-validation approaches, which can lead to an inflation of model performance. In an effort to enhance the generalizability of our models, we used a 10-fold stratified group shuffle split cross-validation approach to ensure that samples from the same participants were not simultaneously distributed across training, validation, and test sets in the same fold. Approximately, 75%, 15%, and 10% of samples were assigned to training, validation, and test sets, respectively.

**Figure 3.**
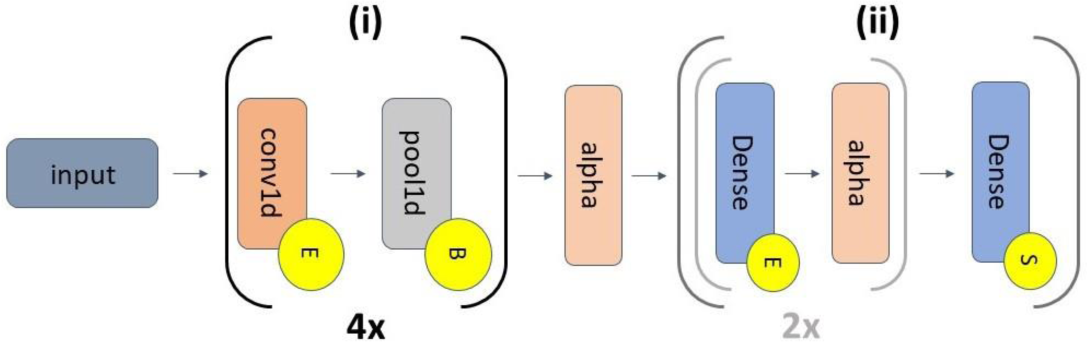
Model Architecture. The model can be subdivided into two segments that are separated by an alpha dropout layer (alpha) – feature extraction (i) and classification (ii). The feature extraction segment repeats 4 times, and the light grey inset within the classification segment repeats twice. Segment (i) has 4 one-dimensional convolutional layers (conv1d) that are each followed by max pooling layers (pool1d). The conv1d layers have 5, 10, 10, and 15 filters and have kernel sizes of 10, 10, 10, and 5. The pool1d layers have pool sizes and strides of 2. Segment (ii) has 3 dense layers with 64, 32, and 2 nodes, respectively. All alpha layers have dropout rates of 0.5. Yellow circles containing an “E”, “B”, or “S” indicate ELU activations, batch normalization, and softmax activations, respectively. Note that conv1d and dense layers have max norm kernel constraints with max values of 1.

During model training, we used a data augmentation approach that has previously been used in [15] to double our training set size. After separating the data into training, validation, and test sets in each fold, we duplicated the training data and augmented the duplicate data via the addition of Gaussian noise (mean = 0, standard deviation = 0.7). We then trained the model on the combined original and augmented data. To account for class imbalances that might randomly occur in the allocation of the training set, we used a class-weighted categorical cross-entropy loss function. We used an Adam optimizer [87] with a learning rate of 0.0075 and a batch size of 128 samples. We trained for a maximum of 35 epochs, using early stopping to end training if validation accuracy (ACC) did not improve after 10 consecutive epochs. To help ensure the generalizability of the model, we also selected the model from the epoch with the peak balanced validation accuracy (BACC). When assessing model test performance, we calculated the mean and standard deviation of the sensitivity (SENS), specificity (SPEC), ACC, and BACC across folds. All convolutional and dense layers, except for the final dense layer which had a softmax activation function with Glorot normal initialization [88], were initialized with He normal initialization [89]. Explainability analyses were performed on the test data from the model with the highest overall BACC.

### Description of Spatial Importance Approach

We applied the αβ-rule [90] of LRP [58], [91] for insight into the relative importance of each channel. LRP is a popular approach in the domain of explainability for image classification and has also been used extensively in the domain of neuroimaging and neurological time-series classification [5], [39], [40], [62], [63], [72], [73], [78], [92]–[97]. We implemented LRP using the Innvestigate library [98]. LRP involves multiple steps. (1) A sample is forward passed through a network. (2) A total relevance value of 1 is assigned to the output node corresponding to the class of interest. (3) The total relevance is iteratively propagated from layer to layer back through the network to the input space using a relevance rule. LRP can propagate both positive (i.e., identifying features that provide evidence for the class of interest) and negative (i.e., identifying features that provide evidence for a class other than the class of interest) relevance. To simplify our analysis by only examining relevance for samples corresponding to their true class, we used the αβ-rule. The αβ-rule has α and β terms, where α and β control positive and negative relevance propagation, respectively. We used α = 1 and β = 0. The equation below shows the αβ-rule.

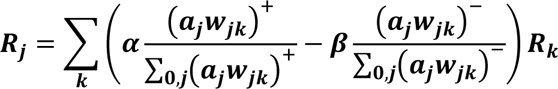

Where the subscripts *k* and *j* correspond to values for one of *K* nodes in a deeper layer and one of *J* nodes in a shallower layer, respectively. The model weights are referenced by *w*, and *a_j_* is the shallower layer activation output.

We output relevance corresponding to the true classes of all test samples in the model with the highest test BACC. While LRP theoretically propagates relevance in a manner that sums to 1, practically, the total relevance can sometimes diverge. As such, after extracting relevance for each test sample, we normalized the absolute relevance of each sample to sum to 100 percent. Specifically, we summed the total absolute relevance assigned to each sample, divided the absolute relevance for each time point and channel by the total absolute relevance, and multiplied by 100. We next summed the total percent of absolute relevance assigned to each channel to estimate spatial importance for each sample. Lastly, to obtain class-specific spatial importance estimates, we averaged separately across HC, MDD, and HC + MDD samples.

The last spatial analysis that we performed sought to determine whether the average spatial importance for each channel was significantly above a uniform spatial distribution of relevance (i.e., where relevance for each channel equals total percent of absolute relevance divided by 19 channels). To this end, we performed a 1-sample t-test comparing the mean relevance of each channel to 100 percent / 19 channels. We then applied false discovery rate (FDR) correction [99] with α = 0.001 to reduce the likelihood of false positive test results. We performed this analysis for HC, MDD, and combined HC and MDD groups.

### Description of Spatial Interaction Approach

After identifying the relative importance of each channel, we sought to understand whether the model uncovered interactions between channels. To this end, we combined spatial LRP as detailed in the previous section with ablation. (1) We output the percent of absolute relevance for each sample and channel *C* (see previous section). (2) We ablated channel *c* of the test samples by replacing it with zeros. While we could have used line-related noise-based ablation approach similar to [63], we elected to use zeros, as line noise was notch filtered during data acquisition. (3) We re-output the percent of absolute relevance for each sample and channel *C*. (4) We calculated the absolute change in relevance belonging to each channel *C*.

The thought process behind our approach was that if a model has uncovered interactions between channels *c*1 and *c*2, then the model should rely upon *c*1 to interpret information in *c*2 and vice versa. Thus, if information in channel *c*1 is removed via ablation and the model uncovered a relationship between channels *c*1 and *c*2, the relevance of channel *c*2 should decrease because the model is no longer able to use the information in channel *c*2 as effectively. Additionally, if the relevance of channel *c*2 increases, then that indicates that the model compensated for the loss of channel *c*1 by relying more upon *c*1. As such, in our approach, we repeated steps 1 through 4 for each of *C* channels and measured the effect of the ablation of channel *c* upon spatial relevance for all of *C* − 1 channels. Our approach relies upon the idea that the relevance of each channel is a combination of its relevance independent of other channels and of its interaction with all other channels.

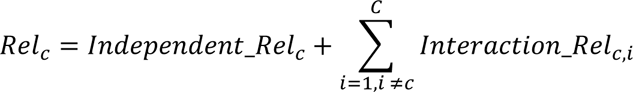

Where *Rel*_*c*_ is the total relevance assigned to channel *c*, *Independent*_*Rel_c_* is the relevance independently assigned to channel *c*, and *Interaction*_*Rel_c,i_* is the relevance of channel *c* that results from interactions between channel *c* and channel *i*, which is not channel *c*.

After outputting the change in relevance of all channels following the ablation of all channels, we sought to determine whether the interactions were statistically significant. To this end, we performed paired two-sample t-tests comparing the relevance assigned to channels before and after the ablation of each channel *c*. We next applied FDR correction [99] with α = 0.001 to reduce the likelihood of false positive test results. We performed this analysis separately for HC, MDD, and combined HC and MDD groups.

### Description of Spatio-Spectral Importance Approach

We next sought to uncover the relative importance of each canonical frequency band in each channel. We analyzed the canonical frequency bands: δ (0 – 4 Hz), θ (4-8 Hz), α (8 – 12 Hz), β (12 – 25 Hz), γ_low_ (25 – 45 Hz), and γ_high_ (55 – 100 Hz). Note that most of γ_high_ was removed during the band pass filtering of the preprocessing, so analyzing γ_high_ enabled us to sanity check our findings, as γ_high_ importance should theoretically be very low. Our spatio-spectral importance analysis consisted of multiple steps. (1) We output the percent of absolute relevance for each test sample and channel *C* (see previous sections). (2) We converted each sample to the frequency domain using a fast Fourier transform (FFT). (3) We assigned coefficients corresponding to frequency band *f* in channel *c* to values of zero. We could have randomly permuted coefficient values or reassigned them from a Gaussian distribution [10], [13], [15], [68]. However, doing so would have required repeatedly perturbing each channel and frequency band dozens of times, which would have been computationally prohibitive given subsequent steps. (4) We re-output the percent of absolute relevance for each sample and channel *C*. (5) We calculated the absolute change in relevance assigned to each channel *c* following the perturbation of frequency band band *f* in channel *c*.

After outputting the change in relevance of all channels following the perturbation of frequency bands, we sought to determine whether the frequency bands in each channel had statistically significant importance. To this end, we performed paired, two-sample, two-tailed t-tests comparing the relevance assigned to channels before and after the ablation of each channel *c*. We next applied FDR correction [99] with α = 0.001 to reduce the likelihood of false positive test results. We performed this analysis separately for HC, MDD, and combined HC and MDD groups.

### Description of Spatio-Spectral Interaction Approach

After identifying the relative importance of each frequency band in each channel, we sought to determine whether the model uncovered interactions between frequency bands *f* in each channel *c* with all other channels *C* not including *c*. This analysis was highly similar to that described in the section, “Description of Spatio-Spectral Importance Approach”. The only difference between the two analyses was that in step 5, we calculated the absolute change in relevance assigned to all channels *C* not including channel *c* following the perturbation of frequency band *f* in channel *c*. After obtaining the percent change in relevance assigned to channels *C* following the the perturbation of frequency band *f* in channel *c*, we employed the same paired, two-tailed, two-sample t-test approach followed by FDR correction described in the previous section.

### Description of Temporal Prototyping and Importance Approach

We lastly sought to uncover any key waveforms differentiating MDDs from HCs. To this end, we combined a prototyping-based approach to identify samples that ideally represented each class and applied LRP to identify the relative importance of each time point and channel for those samples. Our approach consisted of several stages. (1) We input all test samples for both classes into the model and output the activations from the final convolutional layer. (2) We applied principal component analysis (PCA) with 1 PC to reduce the dimensionality of the extracted activations for samples in both classes. (3) We applied k-means clustering to the PC of the activations with 100 initializations sweeping from 2 to 10 clusters. We selected the optimal number of clusters using the maximum silhouette score [100]. We performed clustering separately for each class. (4) We selected the samples closest to the cluster centroids for each class. (5) We output normalized absolute LRP relevance for each sample using the αβ-rule. (6) We applied a moving average with a window size of 20 time points to the relevance assigned to each channel to make visualizing relevance distributions easier. This analysis was two-fold. Firstly, it enabled us to identify samples representative of each class that could be visually inspected for differences. Secondly, it gave insight into how the model analyzed the samples temporally (e.g., Was the relevance temporally distributed or focused on specific highly localized time points? Was the model focused exclusively on unique waveforms or only a few of many similar waveforms?).

## RESULTS

In this section, we describe our model performance, spatial importance, spatial interaction, spatio-spectral importance, spatio-spectral interaction, and temporal prototyping and importance results.

### Model Performance Results

Table 1 shows the classification test performance of our model across all folds. Mean performance for all metrics was above 80%. The model more effectively identified MDDs than HCs, as SENS was at 90% while SPEC was closer to 80%. Additionally, SPEC had a slightly higher standard deviation than SENS. BACC and ACC differed some for specific folds but were on average highly similar. The model test performance for the fold used for explainability was 100% across all metrics.

**Table 1.** Model Performance Results

|  | ACC | BACC | SENS | SPEC |
| --- | --- | --- | --- | --- |
| Mean | 85.00 | 86.52 | 90.33 | 82.70 |
| Standard Deviation | 07.82 | 08.14 | 11.68 | 16.82 |

### Spatial Importance

Figure 4 shows the average total absolute relevance for HCs, MDDs, and both classes combined as well as the t-test results comparing the relevance per channel to a uniform distribution of relevance (i.e., 100% relevance / 19 channels = 5.26% relevance per channel). Across classes, Fp2, F7, Fz, F8, T3, T4, P3, Pz, and P4 were generally below uniform. In contrast, F3, F4, Cz, C4, T5, T6, and O2 generally had above average importance, with F4, T5, and O2 having far above average importance for both classes. Additionally, a few electrodes were significantly above mean importance for one class but not the other. C3 was important for HCs, and Fp1 and O1 were moderately important for MDDs.

**Figure 4.**
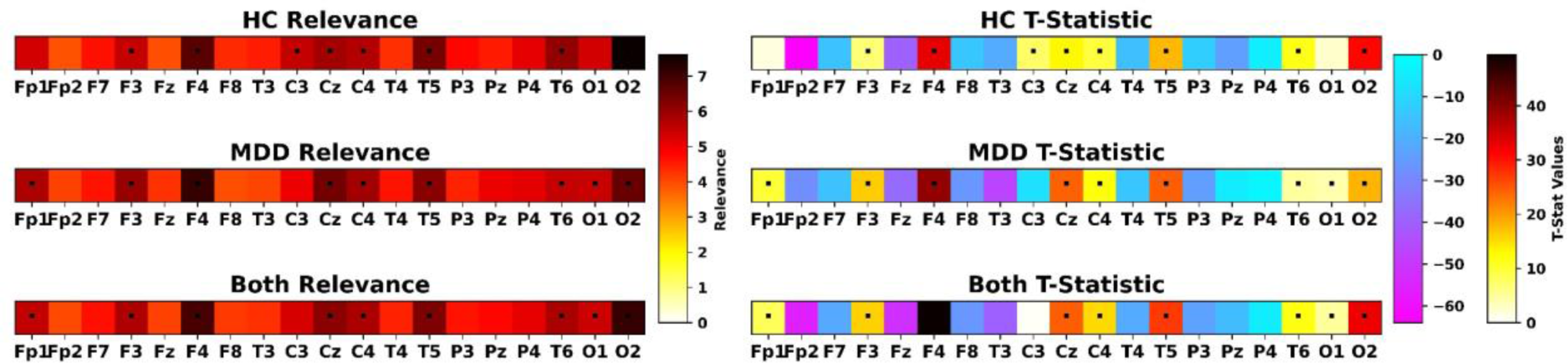
Spatial Importance Results. The leftmost panels show heatmaps of the average relevance for HCs, MDDs, and both classes in descending order. The heatmap to the right of the leftmost panels indicates the amount of relevance corresponding to the heatmap values. The rightmost panels show heatmaps of the t-statistics that resulted from our one-sample, two-tailed t-tests comparing the relevance of each channel to a uniform distribution of relevance (i.e., 100% of relevance / 19 channels). Panels values for HCs, MDDs, and both classes in descending order. Black dots indicate channels with statistically significant p-values following FDR correction (α = 0.001). The two color bars to the right of the leftmost panels indicate the t-statistic values corresponding to the channels with relevance below the uniform distribution and above the uniform distribution. Note that the names corresponding to each channel are displayed along the x-axis.

### Spatial Interactions

Figure 5 shows the results for our spatial interaction analysis. Channels that had a negative change in relevance following the perturbation of another channel can be considered to have an interaction with that channel. A number of channels had a negative t-statistic following the perturbation of other channels, though only a couple of channels had statistically significant reductions in relevance across samples. For HCs, inter-occipital (O1 and O2) interactions were significant. While O1 was not of above-average importance for HCs, O2 was of great importance, and it seems that the model relied upon information in O1 to interpret information in O2. For MDDs, Cz had a significant reduction in relevance following the perturbation of Fp2. Similar to O1 and O2 for HCs, Fp2 was of below average importance for MDDs, but the model seemed to use it to interpret information in Cz, which was highly important for MDDs. Several other frontal, central, and temporal electrodes (F4, F8, C3, and T4) also had reductions in relevance following the perturbation of some frontal (F7, F3, Fz, and F8) and parietal (Pz) electrodes in HCs. While HCs tended to have reductions in more posterior electrodes following the perturbation of other electrodes, MDDs tended to have reductions in more anterior prefrontal, frontal, and central electrode relevance following the perturbation of other frontal, central, and parietal electrodes. While both the HCs and MDDs seemed to have interactions between electrodes, there were few interactions that were identical across classes. Both classes had the same number of significant negative changes in relevance. However, it should be noted that MDDs seemed to have more non-significant negative interactions than HCs.

**Figure 5.**
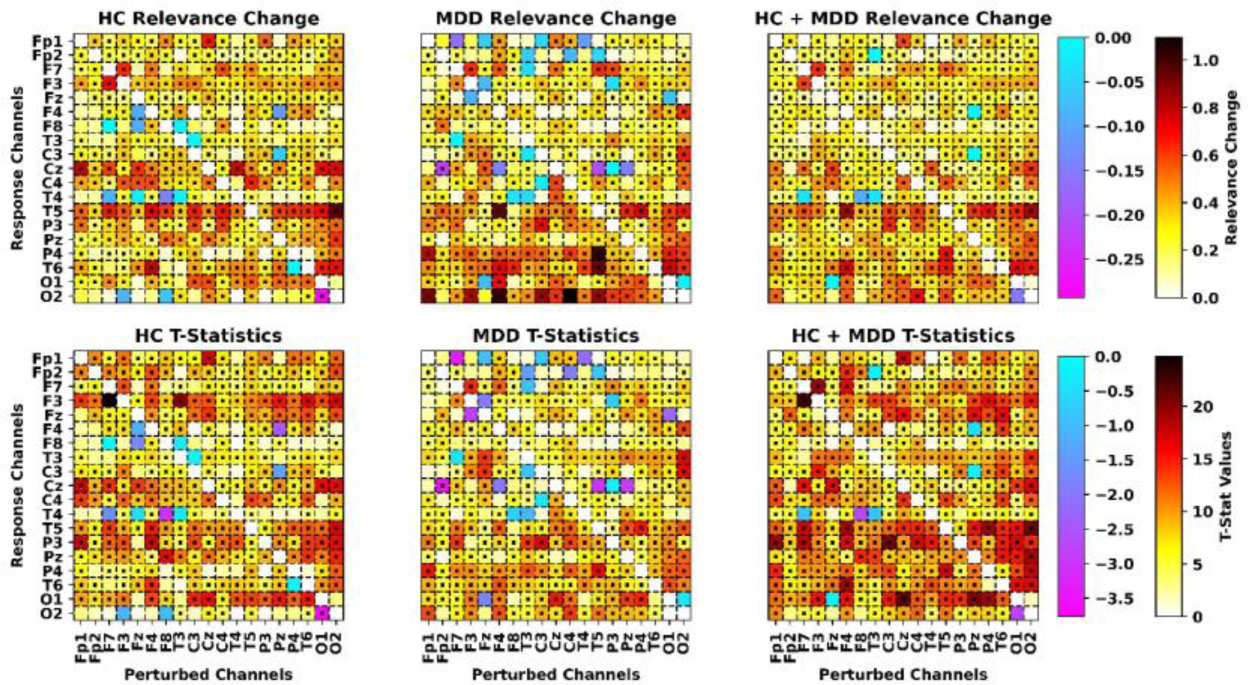
Spatial Interaction Results. From left to right, panels show average relevance interactions for HCs, MDDs, and both classes. The top panels show the change in relevance (relevance 2 – relevance 1), and the bottom panels show t-statistics for the paired, two-tailed, two-sample t-tests comparing the importance of channels before another channel was ablated versus after another channel was ablated. The x-axis indicates channels that were ablated, and the y-axis indicates channels in which a change in relevance is measured (i.e., response channels). Black dots indicate channel combinations in which there was a significant change in the relevance of a response channel after ablation following FDR correction (α = 0.001). Heatmaps to the right of the top and bottom rows of panels indicate the relative magnitude of the change in relevance and the t-statistic values for the t-tests, respectively. Note that values along the left-to-right diagonal were replaced with zeros, as the percent of relevance of a channel decreased much strongly when that channel was itself ablated and such extreme values prevented a visualization of the change in relevance of other channels.

While there were a few channels with significant negative changes in relevance following the perturbation of other channels, a larger number of channels had significant increases in relevance following the perturbation of other channels. In HCs, the model tended to increase relevance to frontal and central electrodes following the ablation of other electrodes. In MDDs, the model tended to increase relevance assigned to temporal electrodes following the ablation of other electrodes. For both classes, the model increased parietal and occipital relevance following the perturbation of other electrodes.

### Spatio-Spectral Importance

Figure 6 shows the importance of each channel and the change in channel relevance following the perturbation of each canonical frequency band. The perturbation of most frequency bands did cause a significant reduction in relevance assigned to their corresponding channels. Nevertheless, there were differences in the magnitude of those effects between HCs and MDDs. The model relied upon θ more strongly and across more channels for identifying HCs than MDDs, and the model relied upon β and γ_low_ across a wider range of channels for identifying MDDs than HCs. Importantly, the model relied upon α for identifying both classes. The model did not rely extensively upon γ_high_. The overall most important frequency and channel combinations for HCs were frontal δ (F3) and θ (F4) and occipital (O2) δ, θ, and α. Overall most important frequency and channel combinations for MDDs were frontal (F3 and F4) α and β, and central (C3, Cz, C4, T4) α and β. Changes in frontal (F7 and F8) and temporal (T4) γ_low_ were also very significant in MDDs.

**Figure 6.**
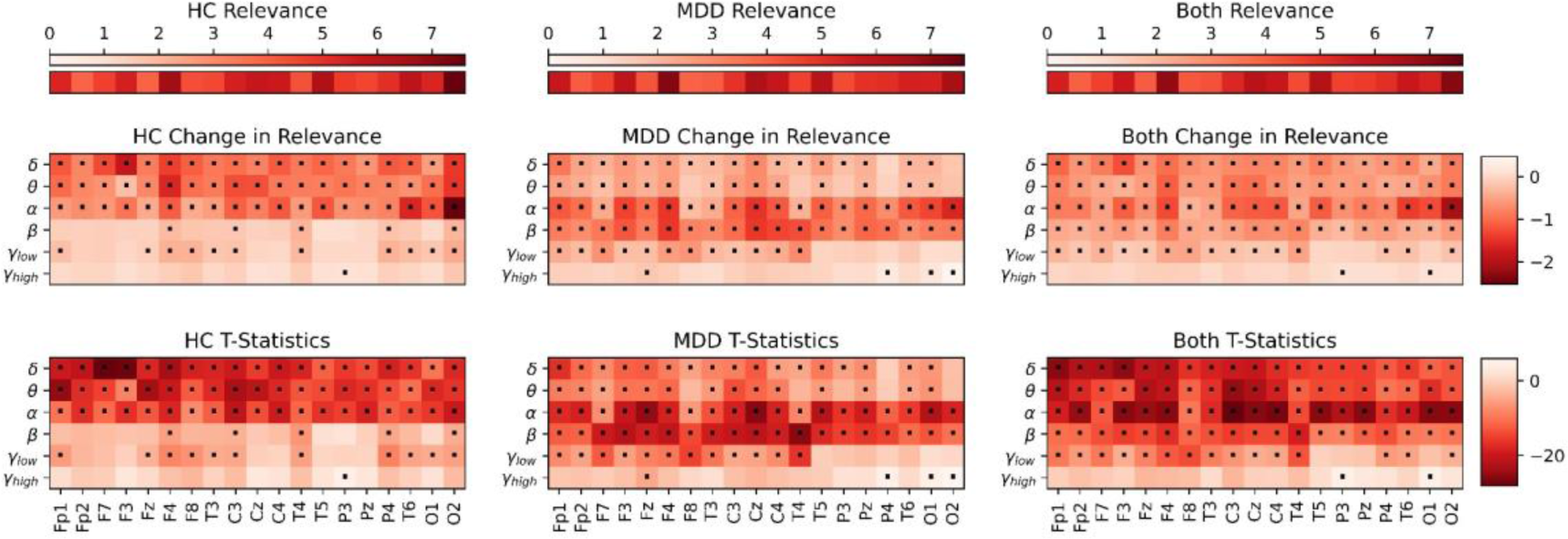
Spatio-Spectral Importance Results. The left, middle, and right columns of panels show results for HCs, MDDs, and both HCs and MDDs, respectively. The top row of panels indicates spatial relevance of each channel. The middle row of panels indicates average change in relevance of specific channels across samples following the perturbation of frequency bands within those channels. The bottom row of panels indicates the t-statistics for the two-sample, two-tailed, paired t-tests comparing the relevance of a channel before versus after perturbation of a frequency band within that channel. Black dots indicate channel and frequency band combinations in which there was a significant reduction in the relevance of a channel after perturbation following FDR correction (α = 0.001). The x-axis shows channels, and the y-axis shows frequency bands. Their corresponding color bars are to the right of the middle and bottom rows, and the corresponding color bars for the top row are located above the panels.

### Spatio-Spectral Interactions

Figure 7 shows spatio-spectral interactions for each frequency band and channel and other channels. Across both classes, spatio-spectral interactions are widespread. As such, we only focus on those interactions that are statistically significant. For HCs, the model also identified some frontal interactions (e.g., F4 with Fp1 θ and F4 and F8 with γ_high_ in a number of channels), central interactions (e.g., C3 with Cz α, T5 and P3 γ_high_, and Pz θ), temporal interactions (e.g., T4 with Fz α, F4 and Cz γ_high_, and P4 and T6 θ), and occipital interactions (e.g., O2 with higher frequency bands in most channels and O1 θ and α). For MDDs, several channels along the prefrontal (Fp1 and FP2), frontal (Fz and F4), and central planes (C3 and Cz) tended to have strong reductions in relevance following the perturbation of most channels. Those channels that did not have negative changes in relevance generally had strongly positive changes. These findings point towards more anterior interactions for MDDs and more posterior interactions for HCs.

**Figure 7.**
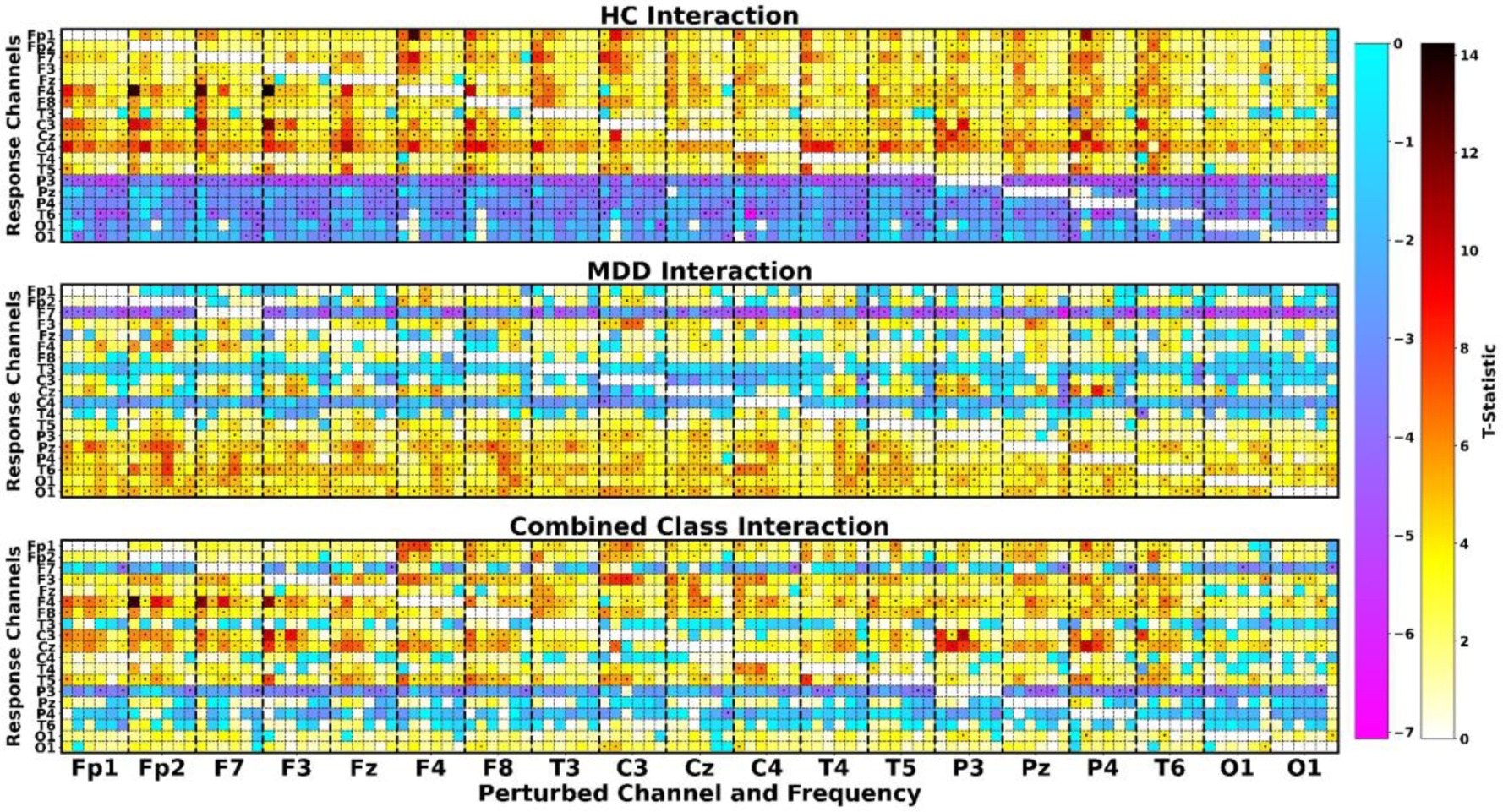
Spatio-Spectral Interaction Results. The top, middle, and bottom panels show heatmaps of the t-statistics from the two-sample, two-tailed, paired t-tests comparing the amount of relevance in a channel before versus after the perturbation of frequency bands in other channels for HCs, MDDs, and both classes, respectively. Perturbed channels and frequency bands are arrayed along the x-axis. Perturbed channels are separated by thick vertical dashed lines, and from left to right within each set of vertical dashed lines are shown results for the perturbation of δ (0 – 4 Hz), θ (4-8 Hz), α (8 – 12 Hz), β (12 – 25 Hz), γlow (25 – 45 Hz), and γhigh (55 – 100 Hz) frequency bands. Channels in which a change in relevance was measured are arrayed along the y-axis (i.e., response channels). The color bars to the right of the figure are shared by all panels and indicate the value of the t-statistics in the heatmaps. Black dots indicate channel and frequency band combinations in which there was a significant change in the relevance of a channel after perturbation following FDR correction (α = 0.001).

### Temporal Prototyping and Importance

Figure 8 shows the extracted CNN features with dimensionality reduced via PCA. It also shows the clusters and samples closest to each cluster centroid. We identified 2 HC clusters and 2 MDD clusters based on the maximal silhouette scores for clustering each class. We used 1 PC that contained approximately 34% of the variance of the extracted CNN features. All other components contained less than 2.5% of the variance. Figure 9 shows the 2 HC and 2 MDD samples closest to each cluster centroid along with an overlayed LRP relevance heatmap that highlights the most important regions of each time-series. MDD and HC clusters were highly separable class-wise. Additionally, while PC 1 had clearly defined MDD1 and MDD2 clusters, HC1 and HC2 were less clearly defined. HC cluster 2 seemed to have a representation more comparable to that of the MDD clusters (i.e., closer to the MDD clusters), and interestingly, the sample for the HC 2 seemed to have more frequent periodic α bursts. While that was the case, HC 1 seemed to have some γ_low_ and MDD 2 had activity at the boundary of β and γ_low_. HC1 also had consistent levels of oscillations at the boundary of δ and θ, which were not present in MDDs. Additionally, both MDDs seemed to contain β oscillations in F7, F8, and T4 that were not as clearly shown in HCs. Both HCs and MDDs seemed to have α oscillations. LRP relevance tended to be more highly concentrated in HCs than MDDs, and MDD 1 relevance was much more concentrated than MDD 2 relevance. In HCs, rather than selecting unique waveforms, the model seemed to focus on a few of many reoccurring waveforms (e.g., α waveforms). This was also often the case for MDD 1, though some unique bursts of higher frequency activity (e.g., between 9 and 10 seconds in Figure 9) were also highlighted. In MDD 2, many periods of α and β (e.g., between 17 and 18 seconds in Figure 9) activity were relevant.

**Figure 8.**
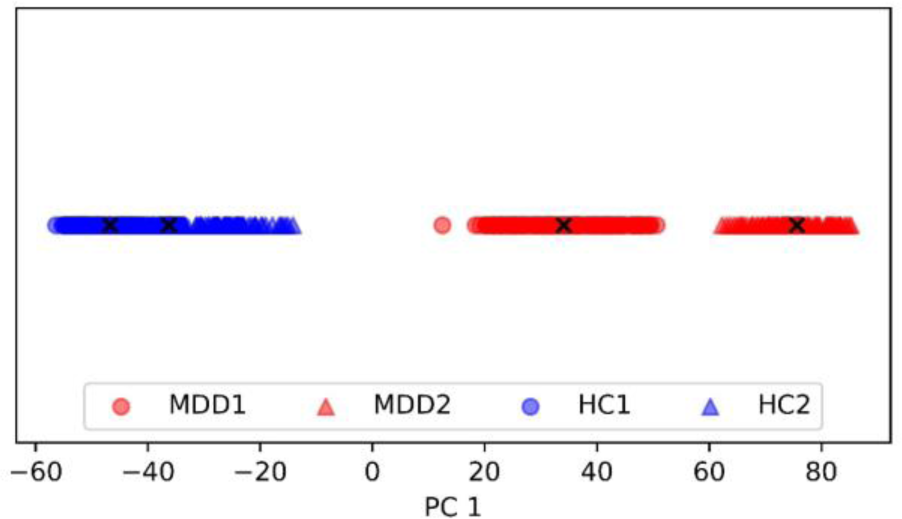
PCA Extracted Activation Clustering Results for Temporal Explainability. One principal component that corresponded to approximately 34% of the variance of the activations for the test samples from the final convolutional layer was used for dimensionality reduction. HC and MDD reduced activations are shown in blue and red, respectively. Two MDD and 2 HC clusters were optimal. Different clusters for each class are each indicated by markers with a different shape, as shown in the legend to the left of the plot. A black “x” is used to mark the samples closest to the cluster centroids that were used in subsequent temporal explainability analyses.

**Figure 9.**
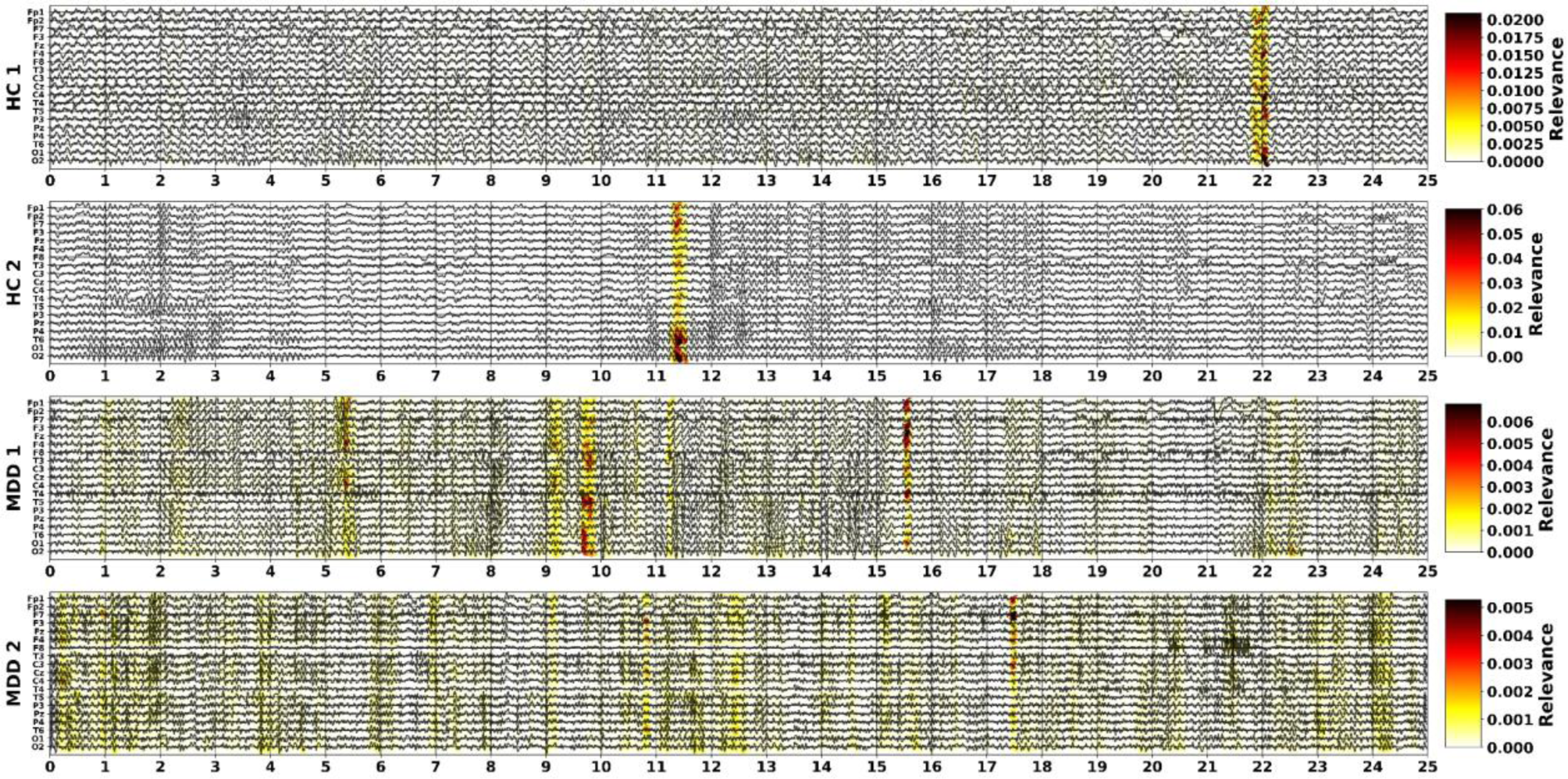
Prototyping and Temporal Importance Results. The time-series of samples closest to the cluster centers of clusters HC 1, HC 2, MDD 1, and MDD 2 (as shown in Figure 8) are displayed in panels from top to bottom. The x-axis shows time in seconds, and the y-axis shows EEG channels. The mean of the data for each channel was subtracted for easier display. A heatmap of LRP importance is overlayed on the time-series. Color bars to the right of the panels show the values of the LRP relevance out of 100% for their respective panels.

## DISCUSSION

In this study, we make two major contributions. (1) We define a taxonomy of explainability approaches for deep learning models trained on raw EEG data. (2) We present a framework for systematically evaluating a deep learning model trained on raw multichannel EEG data that provides insight into each type of feature (i.e., spatial, spectral, and temporal) found in our taxonomy while also expanding upon that taxonomy to provide insight into new types of features (i.e., spatial and spatio-spectral interactions). Importantly, to the best of our knowledge, our methods for examining spatial and spatio-spectral interactions are the first of their kind for raw EEG deep learning classification. Additionally, our approach for spectral importance enables a more sensitive identification of key frequency bands than previous approaches by examining the change in relevance of individual channels (rather than the change in a softmax activation or model accuracy) following perturbation. Both our novel spectral and interaction explainability approaches can provide both local and global insights. Lastly, in contrast to previous approaches that required a specially designed architecture or required that a model be sensitive to the perturbation of input samples, our approach for temporal importance offers an approach for global importance estimation that is applicable to a variety of deep learning classifiers. As a whole in recent years, the field of explainable deep learning for EEG has made great progress, but existing approaches still leave much to be desired. The collection of novel methods presented in our study represents a significant step forward for the field, and the taxonomy that we propose represents a key advancement that will provide guidance for future studies and developments.

### Training a High Performing Model with a Robust Cross-Validation Approach

We developed a model for the classification of individuals with MDD and healthy controls. Our overall model performance was very high (i.e., greater than 80% across all metrics), and the model performance from the fold used in explanations was at 100% across all metrics, increasing the likelihood of the potential generalizability of our explainability findings. Relative to previous studies performing automated diagnosis of MDD using raw EEG data with robust cross-validation techniques, our model obtained higher performance [78], and relative to studies using extracted features with traditional machine learning approaches and robust cross-validation techniques, our model obtained comparable or higher performance [27]. There were some studies that obtained higher test performance than our model using either raw EEG data [80], [81] or extracted features [3], [21], [24], [26]. However, it appears based on the descriptions of their cross-validation approaches that those studies allowed data from the same study participants to leak across training, validation, and test sets within the same folds. That leakage would inflate model test performance and prevent the test performance from actually giving an indication of the generalizability of the patterns learned by their models. This problem in the field is unfortunately relatively common and has been described more extensively in previous studies [27]. Our stratified group shuffle split cross-validation approach protected against this leakage and helped ensure the reliability of our performance findings.

### Identifying Electrodes and Electrode Interactions Important to the Identification of Healthy Individuals and Individuals with MDD

The model relied most upon frontal (F4) and more posterior (T5 and O2) information for identifying both classes indicating that it was able to uncover discriminatory activity for both classes in those areas.

However, the model also relied more strongly upon more frontal (Fp1, F3, F4) and central (Cz) electrodes for identifying MDDs and upon more posterior electrodes for identifying HCs (T6 and O2). This finding is interesting when combined with our channel interaction results. Namely, the model tended to rely upon more frontal and central interactions when identifying MDDs and more posterior interactions when identifying HCs. This finding of spatially widespread effects of MDD is consistent with previous studies that found it necessary to rely upon information from spatially distributed electrodes to obtain high levels of performance [27]. Additionally, it is interesting that while the model relied heavily upon occipital (O1 and O2) electrodes, the model only identified a few interactions between those electrodes and other electrodes across the scalp, and fewer occipital interactions in MDDs than HCs. This could indicate that with eyes closed there are strong effects of MDD on the occipital lobe and that those effects may be related to reduced interactions between occipital areas with frontal areas.

### Examining Why Electrodes are Important by Identifying Their Important Composite Frequency Bands and Frequency Band Interactions

Our findings of spatial importance are further illuminated within the context of our findings on spatio-spectral importance and interactions. While some frequency bands like α that have well-characterized importance in MDD [50], [79] were of widespread importance to the model for identifying both classes, some combinations of frequency bands and channels were important to specific classes. The importance of the posterior electrodes that were important for identifying HCs can be attributed to the presence of θ and α in those electrodes, and the presence of interactions between more posterior electrodes with other electrodes is also found in spatio-spectral interactions where there are widespread interactions with all frequency bands across most channels. The importance of the central electrodes to MDDs is attributable to α and β in those electrodes. While spatial importance indicates that frontal electrodes are important to both classes, spatio-spectral importance indicates that frontal electrodes are important to each class for different reasons. In HCs, frontal δ and θ is more important, and in MDDs, frontal β and γ_low_ are more important. Importantly, frontal θ has been identified as discriminatory between HCs and MDDs [26], and frontal and central β and γ_low_ have previously been associated with inattention in MDD [101]. Additionally, our finding of low γ_high_ importance supports the reliability of the methods, as much of γ_high_ was filtered during initial signal amplification. It is curious that spatio-spectral interactions tend to be more widespread than spatial interactions. This is potentially attributable to how our use of zero-out ablation for identifying spatial interactions and use of spectral perturbation for identifying spatio-spectral interactions interacted differently with the model. Previous studies have shown the importance of choosing ablation and perturbation methods specific to the target domain [63]. While it would have been ideal to be able to use a line noise-related ablation approach in our spatial interactions, the data that we used was publicly available, and line noise was notch filtered during the data collection process. As such, the model would not have learned to consider line noise as neutral information, and the line noise-related ablation approach would thus not be viable.

### Identification of Characteristic Samples for Each Class and Key Waveforms Within Those Samples

Our dimensionality reduction and clustering approach seemed to uncover some underlying structure in the representations learned by the model. For example, from left to right seemed to have longer durations of α activity, and there seemed to be high levels of separation between MDD and HC clusters. Additionally, the model did not seem to uncover unique waveforms of particularly high importance in HCs and MDD 1. Rather it seemed to primarily focus in a highly temporally localized manner on specific waveforms that were identical to many other waveforms found across the 25-second samples. This indicates that while we used 25-second sample sizes, it may have been possible to train an effective classifier with much shorter samples. For MDD 2, the model seemed to have a much more diffuse PCA representation and much more temporally distributed relevance. Additionally, the identified waveforms also illuminate our spatio-spectral importance findings. Specifically, we previously identified that θ was highly important to identifying HCs, and in the samples identified via our prototype approach, HCs had consistently high levels of θ oscillations that were not found in MDDs. Both HCs and MDDs had high levels of α oscillations, which explains why α was important for identifying both classes, and MDDs had high levels of β oscillations, which explains the importance that the model placed upon β for identifying MDDs.

### Limitations and Next Steps

There are several new opportunities for future research directions that are spawned by this study. Our prototyping approach could potentially be expanded upon in future studies. The waveforms identified with our prototyping approach seemed to align well with our previously identified spatio-spectral importance estimates. However, future studies might apply more local explainability approaches to the identified samples to determine how well the findings for the identified prototypes fit with findings for the entire dataset. If there is a high degree of alignment between the findings for each of the prototypes and the global dataset findings, future studies could potentially adapt more robust methods like SHAP [60] for insight into spectral, spatial, and interaction importance that would otherwise not be viable for application with whole EEG datasets given their computational complexity. Additionally, while we applied LRP to provide a measure of channel importance that could change following perturbation and that approach should be broadly applicable to both CNNs and models with recurrent units, future studies might apply approaches similar to ours within the context of other architectures by measuring changes in model attention. Lastly, all of our analyses were performed in the sensor space. Future efforts might use inverse modeling to obtain source space signals and train models on those signals. Resulting explanations could provide enhanced insights into specific brain regions associated with classification performance.

While our proposed explainability approaches were highly effective and present new opportunities for future research, our study methods and findings do have some limitations. Some of the limitations are not unique to our study but rather a problem for the overall field of deep learning-based studies using explainability methods. Specifically, model explanations are not meant to provide an exhaustive investigation of which features could possibly be discriminatory between individuals with MDD and healthy controls. EEG data can be very rich, and as has been shown in previous studies [9], [25], there are often multiple sets of features upon which a model can rely when performing a classification. As such, if our presented methods were to be used in an attempt to obtain exhaustive insight into all of the features that might possibly be useful for diagnosing MDD, it would be necessary to use a more robust training procedure (e.g., many more folds than is the popular practice) and use multiple independently collected datasets. Another limitation of our study findings is related to our dataset size. If we were to try to make generalizable claims about which features are most important for diagnosing MDD, we would need a much larger dataset. Lastly, there are several limitations to the methods proposed in this specific study. Namely, we only perturbed features once, and it would be ideal if we could examine the effects of multiple perturbations upon the same features. Perturbing features only once is relatively common within studies ablating whole channels or modalities, so it is not overly problematic for our spatial interaction analysis. Within spectral importance analyses, it is more common to perturb individual frequency bands more than once; however, due to the computational complexity of repeatedly outputting LRP explanations, we elected to just replace the coefficients of each frequency band with zeros. That said, our use of statistical testing to identify the most important features does help ensure the reliability of our findings. Our use of spectral perturbation may also have caused some edge effects, though this is also a potential problem for all spectral perturbation explainability methods. Alternative approaches might consider applying windows to samples to attenuate any edge effects or performing notch filtering.

## CONCLUSION

The application of deep learning methods to raw EEG data is becoming increasingly common. However, relative to other methods that use traditional machine learning or deep learning with extracted features, deep learning models applied to raw EEG data are less easily explainable. As a result, a field of research has developed seeking to explain these models. In this study, we propose a taxonomy of the explainability methods that have been developed for deep learning models trained on raw EEG. We then introduce an explanatory framework consisting of a series of methods that build upon our proposed taxonomy. In addition to providing insights into key spatial, spectral, and temporal features like existing approaches, the methods in our framework also provide insight into spatial and spatio-spectral interactions uncovered by models. We present our framework within the context of a 1D-CNN trained for automated major depressive disorder diagnosis, identifying interactions between frontal and central electrodes with other electrodes and identifying differences in frontal δ, θ, β, and γ_low_ between healthy individuals and individuals with major depressive disorder. Our study represents a significant step forward for the field of deep learning-based raw EEG classification, providing new capabilities in interaction explainability and providing directions for future research innovations through our proposed taxonomy.

